# Prefrontal cortical pathways mediating cognitive control enhancement from internal capsule stimulation

**DOI:** 10.64898/2026.06.08.730545

**Authors:** Karianne Sretavan, Sumedh S. Nagrale, Victoria Pipia, Elizabeth M. Sachse, Evan M. Dastin-van Rijn, Jaejoong Kim, Spencer Eiting, Aaron N. McInnes, Tara Palnitkar, Jayashree Chandrasekaran, Henry Braun, Samuel Brenny, Yasamin Seddighi, Rémi Patriat, Noam Harel, Matthew D. Johnson, Alik S. Widge

## Abstract

**Background:** Deep brain stimulation (DBS) targeting the ventral internal capsule/ventral striatum (VCVS) is an effective treatment for several psychiatric disorders, but clinical responses vary amongst patients, likely reflecting an incomplete mechanistic understanding of the brain pathways mediating response. VCVS DBS may improve cognitive control, a core deficit in multiple psychiatric disorders. Here, we investigated which prefrontal cortical pathway(s) in the internal capsule are associated with improved cognitive control task performance during VCVS DBS.

**Methods:** Four participants with bilateral VCVS DBS completed the Multi-Source Interference Task (MSIT) across 27 stimulation configurations varying by side (left/right lead), electrode, amplitude, and frequency. Each condition included 100 trials, and response time (RT) assessed task performance. Individualized pathway activation models quantified axonal recruitment across 11 segmented prefrontal cortical fiber bundles per hemisphere. Associations between pathway activation and task RT were analyzed to identify circuits linked to improved cognitive control task performance.

**Results:** Right-sided VCVS stimulation was associated with greater reductions in MSIT RT than left-sided stimulation. Activation of right dorsal prefrontal cortical pathways connected to the dorsolateral prefrontal cortex (dlPFC) and dorsomedial prefrontal cortex (dmPFC) was associated with faster RT, whereas no left-sided pathways were significantly associated with changes in RT.

**Conclusions:** Right-sided dorsal prefrontal pathways connected to the dlPFC and dmPFC are candidate pathways associated with improved cognitive control task performance during VCVS DBS. Pathway activation modeling may inform more precise stimulation strategies for psychiatric disorders characterized by impaired cognitive control, although whether this approach improves clinical outcomes remains to be established.

## INTRODUCTION

Deep brain stimulation (DBS) of the ventral internal capsule/ventral striatum (VCVS) is an FDA-approved treatment for obsessive-compulsive disorder (OCD) (1–3) and under investigation for major depressive disorder (4–6), addiction (7,8), and anorexia (9). While VCVS DBS helps many patients who have not responded to non-surgical treatments, its efficacy remains at 60-70% or lower across indications, limiting clinician and patient willingness to pursue DBS even when indicated (10–12). This is in part due to the field’s incomplete understanding of target engagement, i.e. how to optimize stimulation to engage brain networks mediating response (13,14). These challenges are further exacerbated by a high-dimensional stimulation parameter set that requires tuning of current, frequency, and pulse width in the context of anatomical and clinical heterogeneity (15,16). To date, suggested stimulation targets remain inconsistent (17–21), prompting a need for better target definition and measures of target engagement to improve the effectiveness of DBS for psychiatric applications.

Instead of relying on subjective self-reports, which are often noisy and unreliable (13,14), target engagement might instead be assessed using cognitive constructs that are impaired across psychiatric disorders (14). A particularly relevant construct for VCVS DBS is cognitive control, defined as the ability to flexibly adapt strategies or responses to changing environmental demands (22–26). Deficits in cognitive control are prominent across psychiatric conditions and are thought to drive inflexible behaviors, such as “automatic” negative thoughts in major depressive disorder, habitual drug-taking in addiction even in the absence of reward, and rituals in OCD (27,28). Cognitive control relies on cortico-striato-thalamo-cortical circuitry connecting multiple prefrontal cortex (PFC) subregions to subcortical regions, the same circuitry engaged by VCVS DBS (29–34).

Consistent with that circuit alignment, VCVS DBS improves cognitive control in multiple species. In humans, this manifests as faster response times (RTs) without increased error rates on psychophysical tasks designed to probe cognitive control (26,30,35). The same effect occurs in rats, and computational modeling of both human and rat data has verified that the RT decrease is a true cognitive improvement, not an impulsivity or simple motor effect (36). In both humans and rats, this cognitive effect is dependent on engaging specific VCVS sub-circuits; stimulation in the central to dorsal part of the target appears to produce stronger improvements (30,36). The effect is also lateralized, with larger effects from right-sided stimulation across species (30,37). Finally, rodent evidence suggests that the cognitive effects are mediated by retrograde activation of corticofugal axons passing through the DBS target. Behaviorally effective stimulation was correlated with increased expression of the immediate early gene *c-fos* in rat PFC (36). In a recent optogenetic study, task RTs decreased with optical stimulation of PFC-originating axons but not with striatal grey matter (38).

While these studies demonstrate that stimulating PFC-originating axons in the internal capsule (ICpre) can improve cognitive control, the specific pathway(s) within this broader network mediating this effect remain unclear. ICpre pathways implicated in cognitive control (36,38) are organized topographically: dorsal fibers connect to the dorsolateral and dorsomedial PFC (dlPFC, dmPFC), central fibers to the dorsal anterior cingulate cortex (dACC) and ventrolateral PFC (vlPFC), and ventral fibers to the orbitofrontal cortex (OFC) and ventromedial PFC (vmPFC) (39–41). Despite this organization, it remains unclear which of these specific pathway(s) drives cognitive control improvements, leaving the optimal stimulation target within this region unresolved.

Identifying relevant ICpre pathways could enable personalization of neuromodulatory interventions, including optimization of VCVS DBS to engage individual brain anatomy. In disorders with better-defined therapeutic circuitry, such as Parkinson’s disease and essential tremor, individualized models of pathway activation have been used to guide stimulation toward target networks (42,43), improving surgical targeting precision and post-operative programming effectiveness (44). When target networks are well-characterized, optimization approaches can identify stimulation configurations that maximize engagement of desired pathways while minimizing off-target effects (45). In contrast, for cognitive control, where the relevant circuitry is less well-defined, Bayesian optimization methods applied to behavioral metrics have been used to identify DBS configurations that enhance cognitive control (46,47). Although DBS optimization for psychiatric applications is increasingly feasible, a clearer mechanistic understanding of ICpre pathways underlying cognitive control improvement is necessary to enable more precise, pathway-specific stimulation strategies and improve therapeutic outcomes.

In this study, we investigated which white matter tract(s) within ICpre pathways are associated with improved cognitive control task performance during VCVS DBS. Prior work shows that high-frequency (>100 Hz) stimulation in the central to dorsal portion of these pathways produces stronger cognitive control improvements (30), with effects consistently stronger during right-sided stimulation across studies and species (30,48). Given the topographic organization of ICpre pathways, where dorsal fibers preferentially connect to the dlPFC and dmPFC, these behavioral effects may be associated with engagement of dorsal tracts projecting to these regions, particularly in the right hemisphere. Therefore, we hypothesized that activation of dorsal ICpre tracts, especially those connected to the dlPFC and dmPFC, would be associated with improved cognitive control task performance, with stronger effects during right-sided stimulation.

## METHODS AND MATERIALS

### Participants

Four participants with intractable OCD and bilateral VCVS DBS participated in this study. All implants were placed under a Humanitarian Device Exemption (H050003) or as part of a prior clinical trial (NCT03184454). Two participants received model 3391 DBS leads (Medtronic, Minneapolis, MN, USA), and two received model B33015 SenSight directional leads (Medtronic), connected to model B35200 Percept™ neurostimulators (n=3) or model 37612 Activa™ RC neurostimulators (n=1). All participants were right-handed and completed the task using their right hand. The cohort included three males and one female, aged 21–38 at data collection, with at least 6 months of chronic stimulation exposure. All showed at least partial clinical response to DBS, although this was not fully reflected in Yale-Brown Obsessive-Compulsive Scale (YBOCS) scores for Participant 4. Detailed clinical characteristics are provided in Table 1, and lead locations in Supplementary Figure 1. The sample size was not pre-specified but represents all eligible participants in our geographic region with sufficient high-resolution imaging for modeling.

**Table 1.** Demographics of participants. Participant 4 was able, post-DBS, to complete college and enroll in professional school, to begin driving, and to begin preparing her own food at home, actions which had become impossible due to harm OCD symptoms. However, her YBOCS did not change due to frequent not-just-right bodily experiences and tic-like behaviors that were not changed by DBS.

| Participant | Sex | Age. Yr (at Time of Consent) | Years Since Diagnosis | Pre-Implant YBOCS | Post-Implant YBOCS at Clinically Optimized Setting | DBS Lead |
|---|---|---|---|---|---|---|
| 1 | M | 29 | 17 | 27 | 5 | Medtronic B33015 |
| 2 | M | 38 | 21 | 33 | 23 | Medtronic 3391 |
| 3 | M | 31 | 13 | 28 | 9 | Medtronic B33015 |
| 4 | F | 21 | 4 | 28 | 29 | Medtronic 3391 |

Informed consent was obtained by a study team member not involved in clinical care after full explanation of study procedures and risks. All procedures were approved by the University of Minnesota Institutional Review Board (IRB# 1210M22183, STUDY00021264), and participants provided written informed consent in accordance with the Declaration of Helsinki.

### Behavior Paradigm – Multi-Source Interference Task

To probe cognitive control, participants performed the Multi-Source Interference Task (MSIT; Fig 1). This task has been widely shown to induce statistically robust participant-level effects at both the behavioral and neural level (34,49,50). Importantly, the MSIT has been used extensively in our prior work, establishing it as a sensitive and well-validated readout of DBS engagement of cognitive control circuitry (26,30).

**Figure 1.**
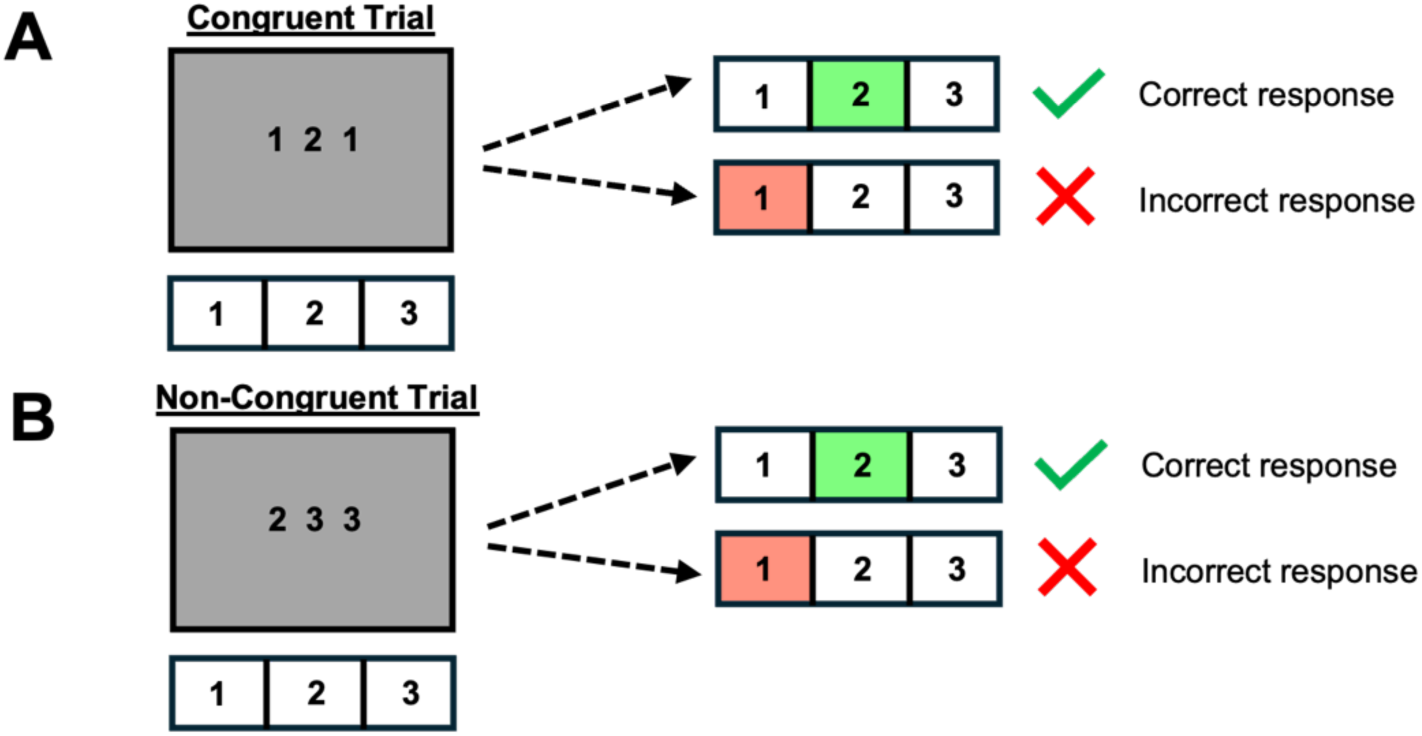
MSIT paradigm showing **(A)** congruent and **(B)** non-congruent trials.

Each trial presented three numbers (ranging 0–3), with two identical digits and one unique digit (the “target”). Participants indicated the identity of the unique number (not its position) via button press. The task included congruent (low-conflict) and non-congruent (high-conflict) trials. In congruent trials, the target’s position matched its correct response location (Fig 1A), whereas in non-congruent trials the target was spatially incongruent (Fig 1B), requiring execution of a non-intuitive visuo-motor mapping (Simon effect), and suppression of distraction from flankers that are also valid targets (Flanker effect). Each block contained 100 trials (50 congruent, 50 non-congruent) presented in pseudo-random order, with constraints preventing more than two consecutive trials of the same interference level or response finger. This design minimized strategy use and increased attentional and cognitive control demands.

Participants completed 27 blocks of the MSIT in one visit; each performed under a single VCVS DBS configuration with one hemisphere stimulated per block. We tested 16 configurations (8 right, 8 left) using 130 Hz stimulation with 150 µs pulse width, varying electrode (0–3, ventral to dorsal) and current amplitude (2 or 4 mA). An additional 8 blocks (4 right, 4 left) used 130 Hz, 210 µs pulse width, with electrodes 0-3 at a fixed 2 mA, enabling comparison of pulse widths at a constant low amplitude for model-based analyses. Three blocks of no stimulation were also included. Configurations were administered in a fixed order across participants due to the small sample size, and potential order effects were assessed separately; see below. Participants were blinded to stimulation condition and on/off status. A complete list of tested stimulation configurations is provided in Supplementary Table 1.

### Imaging

Prior to DBS surgery, all participants underwent pre-operative 7-Tesla (T) MRI at the Center for Magnetic Resonance Research, University of Minnesota, using a Siemens Magnetom Terra scanner (SC72 gradients, 70 mT/m, 100 T/m/s slew rate) with a 32-channel head coil (Nova Medical). The protocol included a 0.6 mm isotropic T1-weighted image, a 1.25 mm isotropic diffusion-weighted scan, and a 0.4 × 0.4 × 1 mm T2-weighted turbo spin-echo image in all participants following previously described methods (51,52). At least 30 days post-surgery, participants received a clinical computed tomography (CT) scan (0.6 mm slices), which was coregistered to the T1 image for DBS lead localization (53,54). T1 images were bias-field corrected (FSL FAST) (55) and skull-stripped (FSL BET) (56). Diffusion data was corrected for motion, eddy currents, and susceptibility distortions using FSL *eddy* and *topup* (57). Structural and diffusion images were then coregistered for subsequent analyses.

### Pathway Activation Modeling Pipeline

Corticofugal, PFC-originating axons are thought to mediate DBS-related improvements in cognitive control (38), motivating our focus on PFC pathways within the internal capsule. To quantify pathway-specific effects of stimulation, we constructed participant-specific computational models combining finite element estimates of stimulation-induced electric fields with biophysical multicompartment axon models to estimate activation of neural pathways near the stimulating electrode (Fig 2). The full pipeline has been previously described in related work (42–44).

**Figure 2.**
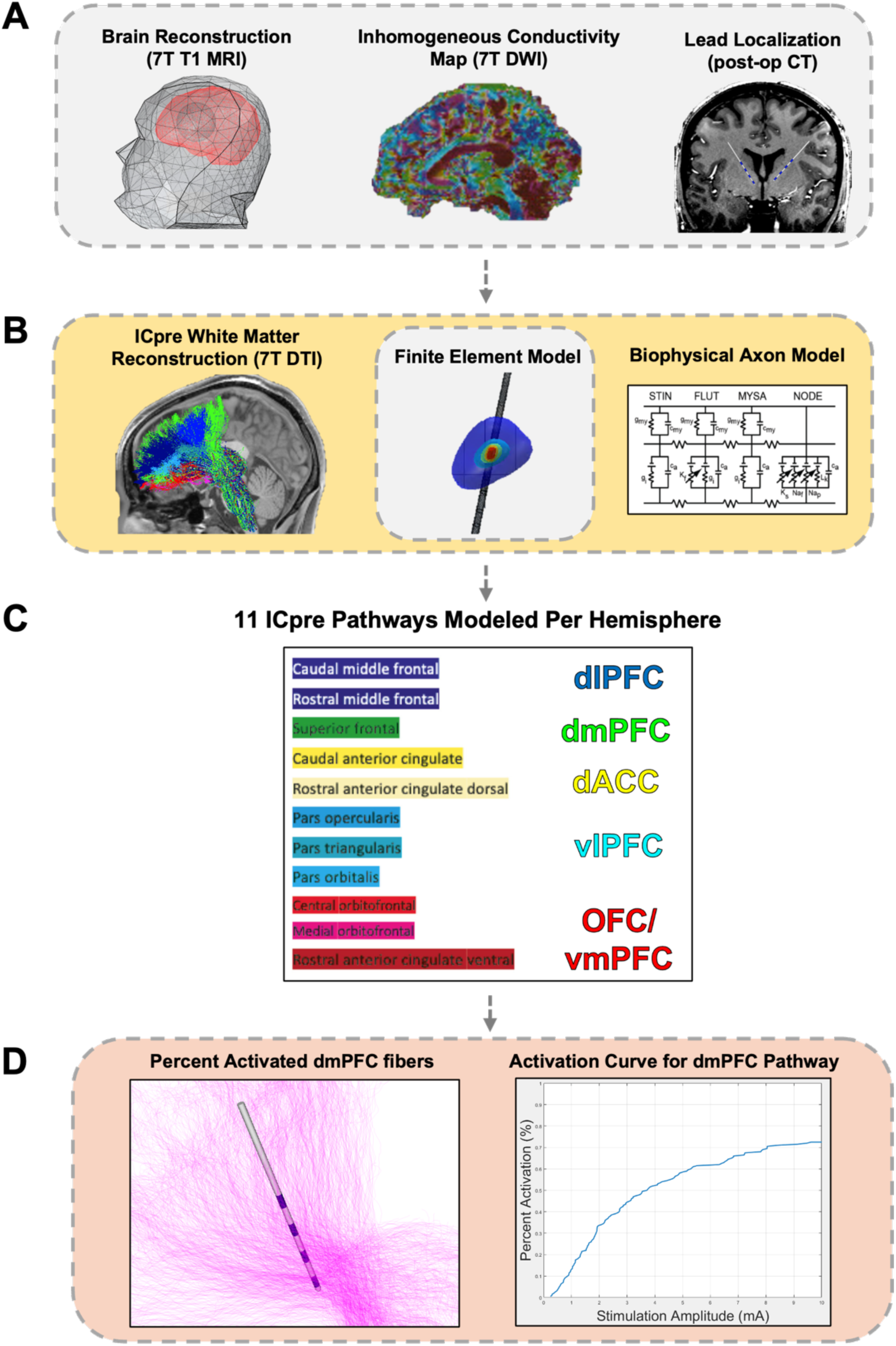
Participant-specific pathway activation modeling pipeline of VCVS DBS. **(A)** Example of a participant’s brain reconstruction from pre-operative MRI and post-operative CT, which informed **(B)** white matter reconstruction of ICpre pathways, a finite element model, and a biophysical multicompartment axon model. **(C)** List of 11 segmented ICpre pathways modeled per hemisphere (left) with corresponding PFC region (right). **(D)** Model-predicted axonal activation of fibers projecting from the left superior frontal gyrus/dmPFC (left, pink) with its corresponding activation curve (right).

Briefly, pre-operative 7T T1-weighted MRI, diffusion-weighted imaging, and post-operative CT were integrated to generate individualized anatomical reconstructions and precisely localize DBS leads. The T1-weighted image defined brain anatomy, diffusion data informed tissue-specific conductivity, and the CT image provided accurate lead placement within the model (Fig 2A). The ICpre pathways were segmented into 22 tracts (Fig 2C) using a validated diffusion tractography algorithm (41), with each tract being defined by its cortical origin and its projection to the thalamus. The pathways were anatomically labeled according to their cortical origins using the Desikan-Killiany atlas. We acknowledge that these anatomical labels do not map directly onto functional cortical regions, which may span multiple gyri and vary across individuals. Corresponding multicompartment axon models were generated for each tract (Fig 2B). Electric field estimates from the finite element model were combined with axon models to compute transmembrane currents and to determine activation thresholds for each axon across pathways and stimulation configurations (Fig 2D).

### Statistical Analysis – Behavior

All statistical analyses were performed in MATLAB (R2022b) and R (R Core Team 2024) using RStudio (version 2025.09.2+418; Posit Team, 2025). The primary behavioral outcome was RT on the MSIT. As participants were pre-trained to very low error rates, accuracy, incorrect, and premature responses (RT < 0.001 s) were not analyzed.

We first evaluated behavioral performance by analyzing trial-level RTs across stimulation conditions (StimOFF, LeftStim, RightStim) and trial types (congruent, non-congruent) using gamma-distributed generalized linear mixed-effects models (GLMs), as used previously (26,30). To assess the effect of stimulation condition on RT, our first model included stimulation condition and trial type as fixed effects and a random intercept for participant to account for repeated measures (RT ∼ StimCondition + TrialType + (1|Participant)). Prior studies have demonstrated that VCVS DBS-related RT reductions occur during both congruent and non-congruent trials (26,30,35,36). Therefore, trial type was included as a covariate to account for systemic differences in RT between trial types when estimating the effect of stimulation condition. Congruent and non-congruent trials were interleaved, which promotes the engagement of proactive cognitive control, which in turn means that stimulation-related changes in RT occur on both trial types, not only on non-congruent trials (36,81). Accordingly, our analysis evaluated stimulation-related changes in RT while accounting for trial type, rather than directly measuring the conflict/interference effect. A post-hoc Wald test of linear contrast (coefTest) was used to compare the fixed effects of LeftStim versus RightStim effects. To assess the effect of trial type on RT, our second model included the StimCondition × TrialType interaction (RT ∼ StimCondition*TrialType + (1|Participant)). Condition-specific post-hoc Wald tests assessed the effect of trial type on RT within each stimulation condition.

We then assessed potential learning or cumulative stimulation effects by examining changes in RT across left and right stimulation blocks (LeftBlock, RightBlock). Trial-level ΔRT was calculated by subtracting the mean RT of the subsequent StimOFF baseline block from each stimulation-trial RT (LeftStim & RightStim). The StimOFF block following all LeftStim blocks was used for LeftStim trials, and the StimOFF block following all RightStim blocks was used for RightStim trials; the first StimOFF block was not used. Positive ΔRT values indicated faster performance during active stimulation relative to baseline, and negative values indicated slower performance during active stimulation relative to baseline. Separate GLMs were fit for left and right stimulation datasets, with block number and trial type as fixed effects and participant as a random intercept (ΔRT ∼ LeftBlocks + TrialType + (1|Participant); ΔRT ∼ RightBlocks + TrialType + (1|Participant)).

To visualize effects of interest (i.e. stimulation condition, trial type, block order), model-adjusted RT and ΔRT values were derived from the GLM outputs. The contribution of all fixed and random effects was removed from the observed RT or ΔRT values, and only the coefficient corresponding to the variable of interest was added back in, isolating its effect on RT or ΔRT for plotting.

### Statistical Analysis – Pathway Activation Models

The next analyses examined the relationship between task performance and pathway activation to identify ICpre pathway(s) associated with cognitive control improvements. A LASSO-penalized generalized linear mixed model (GLMM-LASSO) was used to determine which pathways best predicted ΔRT. This approach combines mixed-effects modeling with L1 regularization, enabling feature selection while accounting for repeated measures. Separate models were fit for left and right stimulation datasets, including block number, trial type, and percent activation of each of the 11 ICpre pathways as fixed effects, with a random intercept for participant (ΔRT ∼ Blocks + TrialType + Pathway 1 + … Pathway 11 + (1|Participant)). The optimal penalty parameter (λ) was selected via leave-one-participant-out cross-validation minimizing mean squared error (MSE). Coefficients at the optimal λ were used to identify pathways most strongly associated with ΔRT. This approach was intended for hypothesis generation rather than generating a predictive clinical model, as a fully held-out test set that would normally be recommended (58) was not feasible given the small sample size.

Using the five pathways identified by GLMM-LASSO, we fit separate unregularized GLMs to evaluate the individual contribution of each pathway. This approach tests whether individual pathways uniquely predict ΔRT while controlling for repeated measures and relevant covariates. Each model included ΔRT as the outcome, with block number, trial type, and percent pathway activation as fixed effects and a participant-level random intercept (ΔRT ∼ Blocks + TrialType + Pathway 1 + (1|Participant)). By examining pathways individually, this method enables us to characterize the direction and magnitude of associations between individual pathway activation and ΔRT. Because the same dataset was used for pathway selection with GLMM-LASSO and subsequent univariate modeling, these analyses were considered exploratory rather than independent validation. Model-adjusted ΔRTs were calculated to characterize the association between each selected pathway and ΔRT while accounting for the included covariates.

To assess whether behavioral performance was better explained by pathway activation or stimulation configuration, we compared mixed-effects models including stimulation configuration alone (ΔRT ∼ TrialType + StimConfiguration + (1|Participant)), stimulation configuration plus pathway activation (ΔRT ∼ TrialType + StimConfiguration + PathwayActivation + (1|Participant)), or pathway activation alone (ΔRT ∼ TrialType + PathwayActivation + (1|Participant)). PathwayActivation includes all 11 ICpre pathways from a single hemisphere. Models were compared using the Akaike Information Criterion (AIC), with lower values indicating better fit.

To determine whether corresponding ICpre pathways exhibited hemispheric differences in modeled activation, we compared the percentage of activated axons during left- and right-sided stimulation across matched stimulation configurations. Comparisons were performed separately for pulse widths of 150 and 210 µs, across configurations varying in electrode and stimulation amplitude. For each configuration, percent activation was compared between hemispheres for each of the 11 corresponding ICpre pathways.

## RESULTS

### Behavior

Four participants with bilateral VCVS DBS implants completed the MSIT under three conditions: StimOFF, LeftStim, and RightStim. We first assessed whether VCVS DBS influenced overall RTs. Both LeftStim and RightStim reduced overall RT relative to StimOFF, with LeftStim on average 29 ms faster (β = −29.45, p = 1.55 x 10^-10^) and RightStim on average 43 ms faster (β = −42.81, p = 4.51 x 10^-21^; Fig 3A). Thus, RT during RightStim was 14 ms faster than LeftStim (β = 14, *F*(1, 10251) = 22.94, p = 1.69 x 10^-6^; Fig. 3A). Participant-level results are shown in Supplementary Figure 5A. Consistent with prior work (49,50), RTs were slower on non-congruent (high conflict) trials versus congruent (low conflict) trials: by 179 ms during StimOFF (β = 179, *F*(1, 10249) = 443.72, p = 1.82 x 10^-96^), 172 ms during LeftStim (β = 172, *F*(1, 10249) = 1688.2, p = 1 x 10^-308^), and 106 ms during RightStim (β = 106, *F*(1, 10249) = 762.49, p = 5.93 x 10^-162^; Fig 3B). Participant-level results are shown in Supplementary Figure 5B. As the order of stimulation configurations was not randomized, potential learning or cumulative effects were examined: ΔRT did not change across blocks during RightStim but was faster by 2.5 ms across LeftStim blocks (β = 2.47, p = 4.15 x 10^-5^; Fig 3C-D). These results replicate our prior reports in independent cohorts (26,30). To further assess potential time-related effects across all stimulation conditions, we examined the association between chronological block order and ΔRT across all 24 active stimulation configurations. Block order showed a modest negative association with ΔRT (β = −0.57, p *=* 0.012; Supplementary Figure 3), indicating that ΔRT was slower by 0.6 ms across all active stimulation blocks.

**Figure 3.**
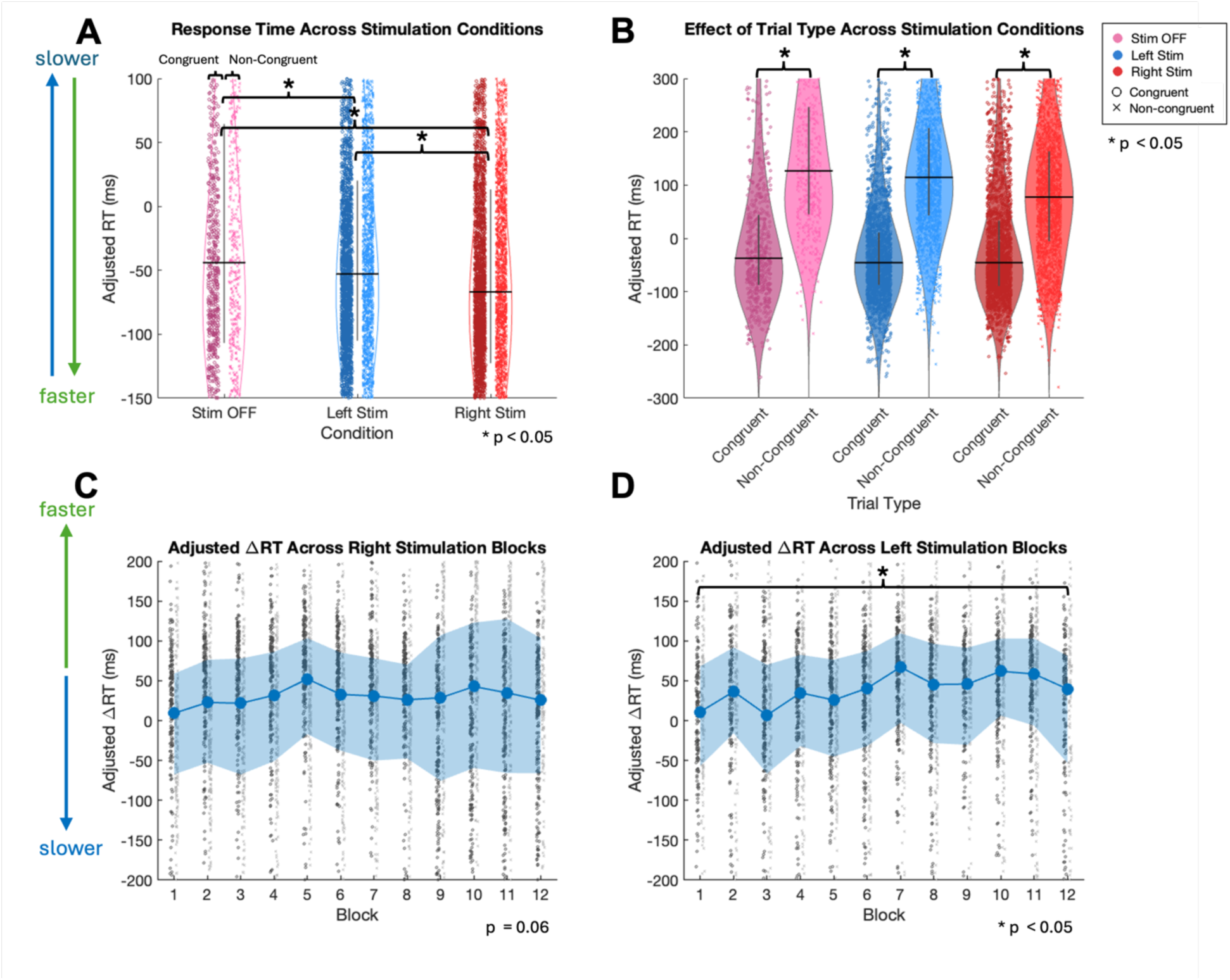
Behavior results from the MSIT. Plots were zoomed to enhance visualization of effects. Complete results showing the full distribution of data points are provided in Supplementary Figure 2. **(A-B)** Violin plots represented distribution of single-trial adjusted RTs (AdjRTs) for each stimulation condition (StimOFF in pink, LeftStim in blue, RightStim in red) with median (black line) and interquartile range (grey bar; 25^th^-75^th^ percentile). Each point corresponded to AdjRT from an individual trial, with dark circles indicating congruent trials (right side) and light crosses indicating non-congruent trials (left side). Participant-level results are shown in Supplementary Figure 5A-B. **(A)** Lower AdjRTs during LeftStim and RightStim compared to StimOFF. Therefore, lower AdjRTs during RightStim compared to LeftStim. **(B)** Higher AdjRTs during non-congruent trials compared to congruent trials during StimOFF, LeftStim, and RightStim. **(C-D)** Adjusted ΔRTs (AdjΔRT) across RightStim and LeftStim blocks with median (blue points) and interquartile range (shaded blue; 25^th^-75^th^ percentile). Each point corresponded to AdjΔRT from an individual trial, with dark circles indicating congruent trials (right side) and light crosses indicating non-congruent trials (left side). A positive value corresponded to a faster AdjΔRT (increased AdjΔRT = faster), and a negative value corresponded to a slower AdjΔRT (decreased AdjΔRT = slower). **(C)** AdjΔRT neither increased nor decreased across RightStim blocks. **(D)** AdjΔRT increased across LeftStim blocks by 2.5 ms.

As all participants were pre-trained and expected to achieve high accuracy, mean accuracy remained high across stimulation conditions (97.2% during StimOFF, 97.8% during LeftStim, and 97.2% during RightStim; Supplementary Figure 4). A binomial logistic mixed-effects model (Accuracy ∼ StimCondition + TrialType + (1|Participant) revealed no significant differences in accuracy between StimOFF and LeftStim (β = 0.25, p = 0.22) or between StimOFF and RightStim (β = −0.02, p = 0.91). The difference between LeftStim and RightStim was also not significant after Bonferroni correction (p = 0.12). Accuracy was significantly lower during non-congruent than congruent trials (β = −0.765, p = 9.81 × 10⁻⁹). These findings provide no evidence of significant stimulation-related differences in accuracy that would suggest a speed-accuracy trade-off.

### Pathway Activation

Our next goal was to relate the behavioral stimulation effects to participant-specific pathway activation models to identify which ICpre pathway(s) are associated with cognitive control improvement. In prior work, stimulation in the central to dorsal sites of the ICpre pathway, particularly in the right hemisphere, produced stronger cognitive control improvements (26,30,48). We therefore hypothesized that activation of right-sided dorsal ICpre pathways connected to dorsal PFC regions would predict faster RTs. In the full model, five right-sided pathways survived LASSO shrinkage. Activation of pathways originating in the caudal middle frontal gyrus (β = 1.96), superior frontal gyrus (β = 4.29), and rostral/dorsal anterior cingulate (β = 1.92) was associated with a faster ΔRT while activation of the caudal anterior cingulate (β = −4.12) and pars orbitalis of the inferior frontal gyrus (β = −1.94) was associated with a slower ΔRT (λ = 6.31, MSE = 0.02; Fig 4A). In contrast, no left-sided pathways survived shrinkage or were associated with ΔRT (λ = 31.62, MSE = 0.02; Fig 4B).

**Figure 4.**
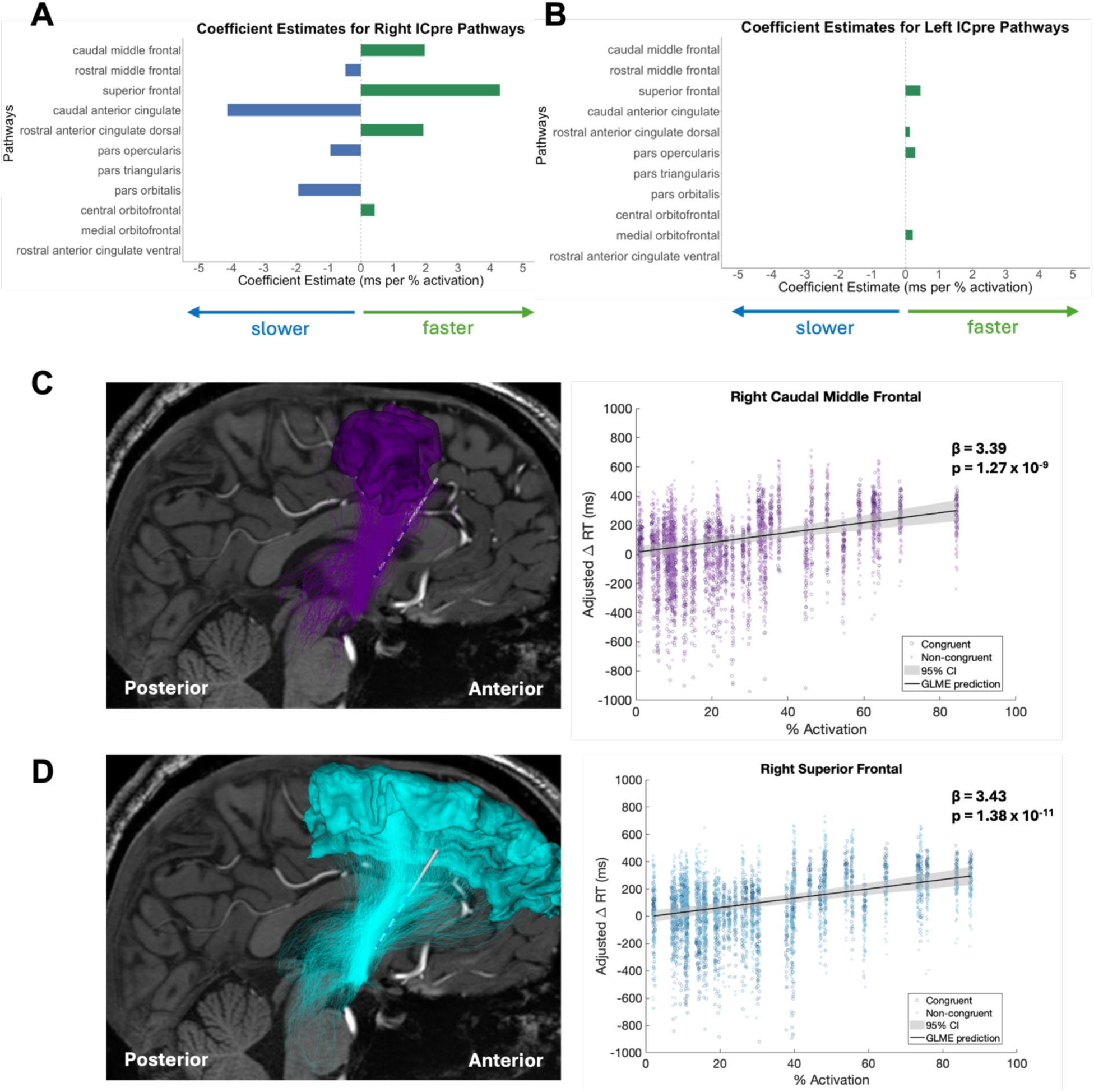
Pathway activation results for significant pathways. **(A-B)** Bar plot showed the fixed-effect coefficients from the GLMM-LASSO at optimal penalty. A positive coefficient reflected pathways associated with faster ΔRT, while a negative coefficient reflected pathways associated with slower ΔRT. **(A)** Three right-sided ICpre pathways predicted faster ΔRT (green bars): caudal middle frontal (β = 1.96), superior frontal, and rostral/dorsal anterior cingulate, while caudal anterior cingulate (β = −4.12) and pars orbitalis predicted slower ΔRT (dark blue bars). **(B)** No left-sided ICpre pathways predicted changes in ΔRT. **(C-D)** Follow-up univariate GLMs assessed the individual predictive contribution of five ICpre pathways identified by the mixed-effects regularized regression and isolated two significant pathways. Left subplot displays example pathways in a sagittal orientation from an individual participant, including the implanted DBS lead and its connectivity to its respective PFC region. Right subplot shows the association between pathway-specific percent activation and adjusted ΔRT across all participants with fixed effects prediction from GLM (solid line) and 95% confidence interval (shaded grey). Each point corresponds to AdjΔRT from an individual trial, with dark circles indicating congruent trials (right side) and light crosses indicating non-congruent trials (left side). Participant-level results are shown in Supplementary Figure 5C-D. Percent activation of tracts connected to the **(C)** right caudal middle frontal gyrus and **(D)** right superior frontal gyrus was associated with faster ΔRT across participants, while the remaining three pathways did not significantly predict either a faster or slower ΔRT. Percent activation values appear discretized due to the use of discrete stimulation amplitudes (2 mA increments), producing step-wise changes in modeled pathway activation. Non-significant results are provided in Supplementary Figure 6.

The mixed-effects regularized regression identified five pathways as potentially predictive of ΔRT. As some of these pathways are highly correlated, LASSO may select multiple pathways that covary with the true drivers of the effect. Therefore, to further characterize the associations of each selected pathway individually, we ran separate unregularized GLMs with participant-specific random intercepts. Because these pathways were selected by GLMM-LASSO using the same dataset, these analyses were considered exploratory and were not treated as independent validation. Of the five selected pathways, only the pathways connected to the right caudal middle frontal gyrus (β = 3.39, p = 1.27 × 10⁻⁹; Fig. 4C) and right superior frontal gyrus (β = 3.43, p = 1.38 × 10⁻¹¹; Fig. 4D) showed significant associations with faster ΔRT. Participant-level results are shown in Supplementary Figure 5C-D. In contrast, the right rostral/dorsal anterior cingulate, caudal anterior cingulate, and pars orbitalis pathways did not show significant associations in the individual models (Supplementary Figure 6).

To determine whether these associations reflected pathway engagement rather than stimulation configuration alone, we compared models including stimulation configuration, pathway activation, or both (Supplementary Table 2). For the right-sided ICpre pathways, inclusion of pathway activation improved model fit relative to stimulation configuration alone (AIC = 11436 vs. 11451.5), and the pathway activation alone model provided the best fit overall (AIC = 11429.7). For the left-side ICpre pathways, the combined pathway activation and stimulation configuration model was the best performing model (AIC = −5381.3) compared to either pathway activation alone (AIC = −5305.1) or stimulation alone models (AIC = −5317.8). In neither hemisphere was a model using stimulation configuration superior. These findings suggest that behavioral performance is more closely related to the degree of pathway engagement than to specific stimulation configurations.

An important consideration in interpreting the right-lateralized behavioral findings is whether hemispheric differences in pathway recruitment contribute to this asymmetry (Supplementary Figure 7). Comparing modeled activation of ICpre pathways during left- and right-sided stimulation revealed configuration-dependent differences in pathway recruitment. At the most ventral electrode (contact 0), activation was higher during left-sided stimulation across all pathways at both pulse widths (150 and 210 μs), whereas more dorsal configurations (contacts 1-3) showed generally higher activation during right-sided stimulation.

## DISCUSSION

We investigated which ICpre pathway(s) were associated with enhanced cognitive control task performance during VCVS DBS. We identified dorsal ICpre pathways connected to the right caudal middle frontal gyrus, part of the dlPFC, and the right superior frontal gyrus, part of the dmPFC, as associated with improved MSIT performance (faster response times). In contrast, no left-sided ICpre pathways were associated with changes in MSIT response times. These findings suggest that engagement of right-sided dorsal ICpre pathways connected to the dlPFC and dmPFC may contribute to improved cognitive control task performance during VCVS DBS. Identifying candidate pathways associated with these behavioral effects may inform the development of more precise, mechanistically guided DBS programming strategies by helping optimize stimulation parameters (including electrode configuration, current amplitude, frequency, and pulse width) to preferentially engage pathways associated with improved performance, while minimizing engagement of other pathways that may be counter-productive. Whether selectively engaging these pathways can improve psychiatric symptoms, however, remains to be established.

### Dorsal PFC

The dlPFC and dmPFC jointly support cognitive control through a monitoring-regulation framework in which the dmPFC detects conflict and signals the need for increased control, recruiting the dlPFC to implement task-appropriate adjustments (59–61). These regions are structurally connected along an anterior-posterior gradient, with posterior dmPFC connected to posterior dlPFC and anterior dmPFC connected to anterior dlPFC (62). In the context of our findings, this functional and structural connectivity provides a potential mechanistic account of successful MSIT performance. In particular, coordinated dmPFC-dlPFC activity may enable monitoring signals to drive flexible control adjustments in the presence of competing rule demands across congruent and non-congruent trials.

Disruption of dorsal ICpre pathways is widely implicated in psychiatric disorders. Both the dlPFC and dmPFC are key nodes in cortico-striatal-thalamo-cortical circuitry, which is frequently altered in OCD, major depressive disorder, and anxiety disorders (63–65). Converging evidence from VCVS DBS studies further highlight the importance of dlPFC-associated pathways, with circuit engagement predicting therapeutic outcomes (17,19).

Together, these findings indicate that dmPFC-dlPFC interactions are central to cognitive control and may represent a substrate of psychiatric dysfunction. Targeting ICpre pathways connected to these regions via VCVS DBS may provide a strategy for modulating cognitive control networks. Whether such modulation translates to clinical improvement, and whether their effects generalize across psychiatric disorders, remain important questions for future investigation.

### VCVS DBS Laterality Effects

MSIT RT improvement was associated with engagement of right dorsal ICpre projections from the dlPFC and dmPFC, while no left-sided pathways were associated. This asymmetry, favoring right-sided stimulation, has been previously reported in human and animal models (30,48) and challenges the assumption that bilateral VCVS DBS is necessary to achieve optimal therapeutic outcomes. Prior work suggests that for some patients, unilateral DBS may be sufficient to alleviate symptoms (30,48), potentially achieving similar benefits as bilateral stimulation (66,67). This approach could potentially reduce surgical time and the duration required to optimize parameters for each patient.

Right-lateralized effects may reflect asymmetries in structural connectivity and functional organization of cognitive control systems. Interhemispheric prefrontal connectivity provides one potential mechanism, as prefrontal regions are interconnected across hemispheres and project bilaterally to the striatum and other basal ganglia structures (68), allowing unilateral stimulation to distribute modulatory effects across both hemispheres. Within this framework, the right dlPFC and dmPFC may function as dominant “control hubs”, where right-sided stimulation may directly engage these circuits, while left-sided stimulation may act indirectly via interhemispheric pathways (e.g., corpus callosum). Such indirect signaling may reduce the efficacy or precision of network engagement, yielding weaker behavioral effects.

Task demands may further explain this asymmetry. In the Dual Mechanisms of Control framework, cognitive control operates through proactive control, which involves sustained maintenance of goal-relevant information, and reactive control, which is recruited following stimulus onset for conflict resolution (69–73). While the dlPFC supports both processes (71,74), proactive control is proposed to be more left-lateralized (71,75,76), and reactive control more right-lateralized (75). Our MSIT design unpredictably intermixes congruent and non-congruent trials. This minimizes the effectiveness of anticipatory strategies and emphasizes reactive control. Accordingly, right-sided VCVS stimulation may preferentially engage right ICpre circuits, including dlPFC-dmPFC pathways supporting rapid conflict detection and resolution, whereas left-lateralized proactive systems are less engaged. This laterality effect therefore likely reflects selective modulation of right-hemispheric reactive control networks aligned with task demands.

The observed hemispheric differences in modeled pathway activation (Supplementary Figure 7) suggest that differences in tract geometry, electrode placement, and pathway recruitment may contribute to the right-lateralized behavioural findings. However, our analysis characterizing hemispheric differences did not establish whether these differences explain the observed behavioral asymmetry. As stimulation configurations varied systematically in electrode and amplitude, we could not isolate the contribution of pathway recruitment to the behavioral findings. Additionally, inherent asymmetries in the anatomical organization of left versus right ICpre pathways, as well as functional hemispheric specialization, may also contribute to the observed asymmetry, but these possibilities could not be directly assessed in the present study. Future studies using systematically matched stimulation configurations and behavioral assessments are needed to disentangle these contributions.

While our findings are specific to VCVS DBS, the potential relevance of stimulation laterality may extend to other neuromodulation approaches, including transcranial magnetic stimulation. Unilateral dlPFC has demonstrated antidepressant efficacy, though randomized trials and meta-analyses have not consistently shown bilateral stimulation to be superior to unilateral stimulation (77–80). These findings suggest that unilateral stimulation could be effective in psychiatric neuromodulation, although differences between TMS and VCVS DBS limit direct comparison. Further research is needed to determine whether the right-lateralized associations with cognitive control task performance observed in this study translate to clinical benefits from unilateral VCVS DBS.

### Limitations and Future Directions

The small sample size of four participants is an important limitation, reflecting the rarity of individuals with both VCVS DBS implants and high-resolution imaging. To mitigate this constraint, each participant completed a large number of MSIT trials across 27 stimulation configurations (12 LeftStim, 12 RightStim, 3 StimOFF), with 100 trials per setting. This intensive within-subject design strengthens comparisons across stimulation configurations, but larger and more diverse cohorts are necessary to improve generalizability.

Because this study focused on unilateral right- and left-sided VCVS stimulation, we could not directly evaluate the additive or synergistic effects of bilateral stimulation. Although prior work suggests that bilateral stimulation may not provide additional cognitive benefits beyond unilateral effects (48), our findings do not establish whether unilateral stimulation is sufficient to achieve optimal therapeutic outcomes or whether bilateral stimulation provides additional clinical benefit. Direct comparison between the two in humans remains necessary to investigate.

The fixed order of stimulation configurations presents a potential source of bias, as fatigue or stimulation carryover could influence task performance over time. To assess potential time-related effects, we examined the association between chronological block order and ΔRT across all 24 active stimulation configurations. Block order was associated with a decrease in ΔRT, indicating a slower RT over the session. This relationship was opposite to the main finding that activation of right-sided PFC pathways was associated with faster RT, arguing against a practice effect as an explanation for the main finding. Furthermore, prior human experiments using the same task and similar stimulation paradigms reported an absence of practice effects (26,30), further supporting the interpretation that practice effects are unlikely to explain the present findings. Nevertheless, because stimulation side, pulse width, electrode, and amplitude varied systematically with order, other order-related effects cannot be excluded. Future studies should randomize or counterbalance stimulation configurations to better distinguish stimulation-specific effects from time-dependent changes in task performance.

Our computational modeling approach leverages individualized imaging, including high-resolution DTI-derived structural pathways and tissue conductivity properties accounting for inhomogeneity and anisotropy, to capture individual neuroanatomical variability. However, these models rely on simplifying assumptions and cannot fully replicate *in vivo* conditions. Key limitations include uncertainties in diffusion tractography (e.g., crossing fibers), literature-derived conductivity estimates, simplified tissue segmentation, and uniform axon diameters. While these factors may affect predicted activation patterns, their precise impact is unknown, highlighting the need to determine the model complexity required to accurately represent neural dynamics. Additionally, we acknowledge that the Desikan-Killiany atlas labels used to identify cortical regions of origin, such as the superior frontal and caudal middle frontal gyri, do not map exactly onto the broader functional regions of the dmPFC and dlPFC, respectively. These functional regions span multiple gyri, and their anatomical boundaries may vary across individuals, which should be considered when interpreting the pathway labels.

Establishing the validity of predicted pathway engagement also remains an important challenge. Non-invasive electrophysiological recordings such as electroencephalography could assess whether VCVS DBS targeting the right dlPFC-dmPFC ICpre pathways provides corresponding cortical activity consistent with improved cognitive control. However, it cannot directly confirm specific pathway engagement. Alternatively, animal models provide a more direct approach; homologous rodent VCVS DBS models improve cognitive control (36,48), and when combined with optogenetics, could enable selective manipulation of medial and lateral PFC projections that are partly homologous to the human dlPFC and dmPFC ICpre pathways, allowing direct validation of DBS effects on specific ICpre pathways (38).

Lastly, the present study assessed cognitive control task performance rather than directly assessing clinical therapeutic outcomes. Although the identified pathway-behavior associations may help characterize cognitive control network engagement during VCVS DBS, we did not directly assess whether recruitment of these pathways was associated with improvement in OCD symptoms or other psychiatric outcomes. Recent work suggests potential links between cognitive control and major depression symptoms (81); however, whether enhancing cognitive control translates to clinical benefit, and whether such benefits generalize across psychiatric disorders, remains unclear. Therefore, the identified pathways in this study should be considered candidates for further investigation. Future studies integrating pathway-specific activation modeling, cognitive control measures, and longitudinal clinical outcomes will be needed to determine their relevance to therapeutic programming.

## CONCLUSIONS

DBS targeting the VCVS enhances cognitive control by stimulating PFC-originating axons in the internal capsule. However, the specific ICpre pathways underlying these behavioral effects have remained unresolved. In this study, we generated individualized models of neural activation across multiple stimulation configurations during completion of the MSIT to identify the ICpre pathways associated with improved cognitive control task performance. Our findings demonstrate that right-sided VCVS DBS was associated with greater improvements in MSIT response time than left-sided stimulation. Moreover, this behavioral effect is supported by pathway activation results, which show that activation of right-sided dorsal ICpre pathways connected to the dlPFC and dmPFC was associated with faster response times, whereas no left-sided ICpre pathways were associated with changes in response time. These findings identify candidate pathways for further investigation and suggest that optimizing VCVS DBS target and programming to preferentially engage right-sided dorsal ICpre pathways connected to the dlPFC and dmPFC may provide a basis for more precise, mechanistically guided treatment strategies. Whether this approach improves therapeutic outcomes for psychiatric disorders characterized by impaired cognitive control remains to be established.

## DATA AVAILABILITY STATEMENT

The data are available from the corresponding author on reasonable request, subject to applicable ethical and legal restrictions.

## AUTHOR CONTRIBUTION

Conception and design of work: KS, NH, MDJ, ASW. Data acquisition: SSN, VP, EMDR, JK, SE, ANM, TP, JC, HB, SB, YS, RP. Data analysis: KS, EMS, NH, MDJ, ASW. Data interpretation: KS, NH, MDJ, ASW. Drafting manuscript: KS. Revising manuscript: KS, TP, JC, HB, SB, YS, RP, NH, MDJ, ASW. Final approval: KS, SSN, VP, EMS, EMDR, JK, SE, ANM, TP, JC, HB, SB, YS, RP, NH, MFJ, ASW.

## LEGEND FOR FIGURES

**Figure 1**. MSIT paradigm showing **(A)** congruent and **(B)** non-congruent trials.

**Figure 2**. Participant-specific pathway activation modeling pipeline of VCVS DBS. **(A)** Example of a participant’s brain reconstruction from pre-operative MRI and post-operative CT, which informed **(B)** white matter reconstruction of ICpre pathways, a finite element model, and a biophysical multicompartment axon model. **(C)** List of 11 segmented ICpre pathways modeled per hemisphere (left) with corresponding PFC region (right). **(D)** Model-predicted axonal activation of fibers projecting from the left superior frontal gyrus/dmPFC (left, pink) with its corresponding activation curve (right).

**Figure 3**. Behavior results from the MSIT. Plots were zoomed to enhance visualization of effects. Complete results showing the full distribution of data points are provided in Supplementary Figure 2. **(A-B)** Violin plots represented distribution of single-trial adjusted RTs (AdjRTs) for each stimulation condition (StimOFF in pink, LeftStim in blue, RightStim in red) with median (black line) and interquartile range (grey bar; 25^th^-75^th^ percentile). Each point corresponded to AdjRT from an individual trial, with dark circles indicating congruent trials (right side) and light crosses indicating non-congruent trials (left side). Participant-level results are shown in Supplementary Figure 5A-B. **(A)** Lower AdjRTs during LeftStim and RightStim compared to StimOFF. Therefore, lower AdjRTs during RightStim compared to LeftStim. **(B)** Higher AdjRTs during non-congruent trials compared to congruent trials during StimOFF, LeftStim, and RightStim. **(C-D)** Adjusted ΔRTs (AdjΔRT) across RightStim and LeftStim blocks with median (blue points) and interquartile range (shaded blue; 25^th^-75^th^ percentile). Each point corresponded to AdjΔRT from an individual trial, with dark circles indicating congruent trials (right side) and light crosses indicating non-congruent trials (left side). A positive value corresponded to a faster AdjΔRT (increased AdjΔRT = faster), and a negative value corresponded to a slower AdjΔRT (decreased AdjΔRT = slower). **(C)** AdjΔRT neither increased nor decreased across RightStim blocks. **(D)** AdjΔRT increased across LeftStim blocks by 2.5 ms.

**Figure 4**. Pathway activation results for significant pathways. **(A-B)** Bar plot showed the fixed-effect coefficients from the GLMM-LASSO at optimal penalty. A positive coefficient reflected pathways associated with faster ΔRT, while a negative coefficient reflected pathways associated with slower ΔRT. **(A)** Three right-sided ICpre pathways predicted faster ΔRT (green bars): caudal middle frontal (β = 1.96), superior frontal, and rostral/dorsal anterior cingulate, while caudal anterior cingulate (β = −4.12) and pars orbitalis predicted slower ΔRT (dark blue bars). **(B)** No left-sided ICpre pathways predicted changes in ΔRT. **(C-D)** Follow-up univariate GLMs assessed the individual predictive contribution of five ICpre pathways identified by the mixed-effects regularized regression and isolated two significant pathways. Left subplot displays example pathways in a sagittal orientation from an individual participant, including the implanted DBS lead and its connectivity to its respective PFC region. Right subplot shows the association between pathway-specific percent activation and adjusted ΔRT across all participants with fixed effects prediction from GLM (solid line) and 95% confidence interval (shaded grey). Each point corresponds to AdjΔRT from an individual trial, with dark circles indicating congruent trials (right side) and light crosses indicating non-congruent trials (left side). Participant-level results are shown in Supplementary Figure 5C-D. Percent activation of tracts connected to the **(C)** right caudal middle frontal gyrus and **(D)** right superior frontal gyrus was associated with faster ΔRT across participants, while the remaining three pathways did not significantly predict either a faster or slower ΔRT. Percent activation values appear discretized due to the use of discrete stimulation amplitudes (2 mA increments), producing step-wise changes in modeled pathway activation. Non-significant results are provided in Supplementary Figure 6.

