## Supplemental figures and tables for "Prefrontal cortical pathways mediating cognitive control enhancement from internal capsule stimulation"

**SUPPLEMENTAL INFORMATION**

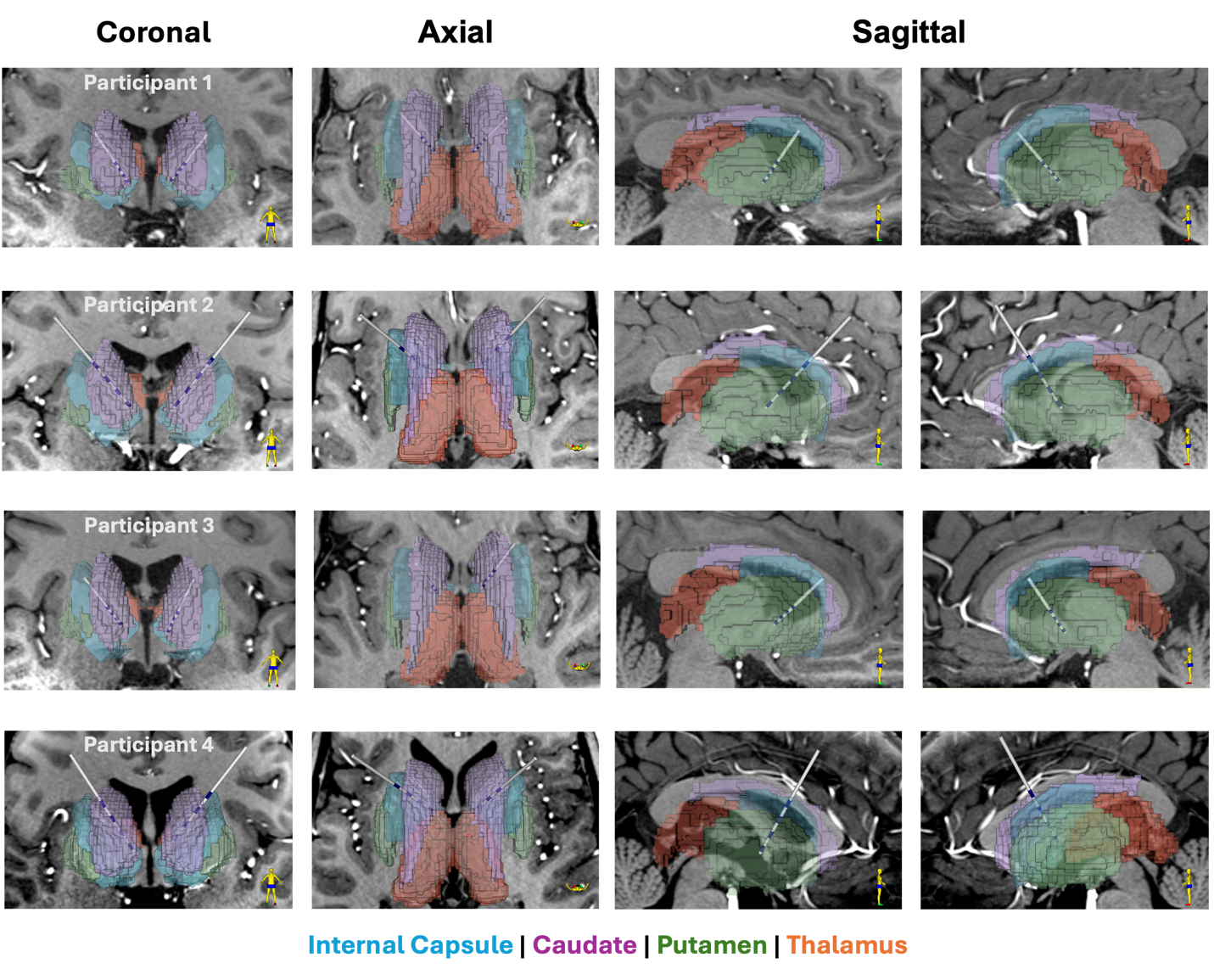

**Supplementary Figure 1.** DBS lead location for all study participants with 7T T1-weighted background image and participant-specific atlas-based segmentations of surrounding grey matter structures using the FreeSurfer Desikan-Killiany atlas in native T1 space (77) (internal capsule = blue, caudate = purple, putamen = green, thalamus = orange).

| **Condition** | **Electrode (0-3)** | **Amplitude (mA)** | **Pulse With (μs)** | **Frequency (Hz)** |
| --- | --- | --- | --- | --- |
| DBS OFF | N/A | N/A | N/A | N/A |
| LEFT | 0 | 2 mA | 150 μs | 130 Hz |
| LEFT | 0 | 4 mA | 150 μs | 130 Hz |
| LEFT | 1 | 2 mA | 150 μs | 130 Hz |
| LEFT | 1 | 4 mA | 150 μs | 130 Hz |
| LEFT | 2 | 2 mA | 150 μs | 130 Hz |
| LEFT | 2 | 4 mA | 150 μs | 130 Hz |
| LEFT | 3 | 2 mA | 150 μs | 130 Hz |
| LEFT | 3 | 4 mA | 150 μs | 130 Hz |
| DBS OFF | N/A | N/A | N/A | N/A |
| RIGHT | 0 | 2 mA | 150 μs | 130 Hz |
| RIGHT | 0 | 4 mA | 150 μs | 130 Hz |
| RIGHT | 1 | 2 mA | 150 μs | 130 Hz |
| RIGHT | 1 | 4 mA | 150 μs | 130 Hz |
| RIGHT | 2 | 2 mA | 150 μs | 130 Hz |
| RIGHT | 2 | 4 mA | 150 μs | 130 Hz |
| RIGHT | 3 | 2 mA | 150 μs | 130 Hz |
| RIGHT | 3 | 4 mA | 150 μs | 130 Hz |
| RIGHT | 0 | 2 mA | 210 μs | 130 Hz |
| RIGHT | 1 | 2 mA | 210 μs | 130 Hz |
| RIGHT | 2 | 2 mA | 210 μs | 130 Hz |
| RIGHT | 3 | 2 mA | 210 μs | 130 Hz |
| LEFT | 0 | 2 mA | 210 μs | 130 Hz |
| LEFT | 1 | 2 mA | 210 μs | 130 Hz |
| LEFT | 2 | 2 mA | 210 μs | 130 Hz |
| LEFT | 3 | 2 mA | 210 μs | 130 Hz |
| DBS OFF | N/A | N/A | N/A | N/A |

**Supplementary Table 1.** List of tested stimulation configurations.

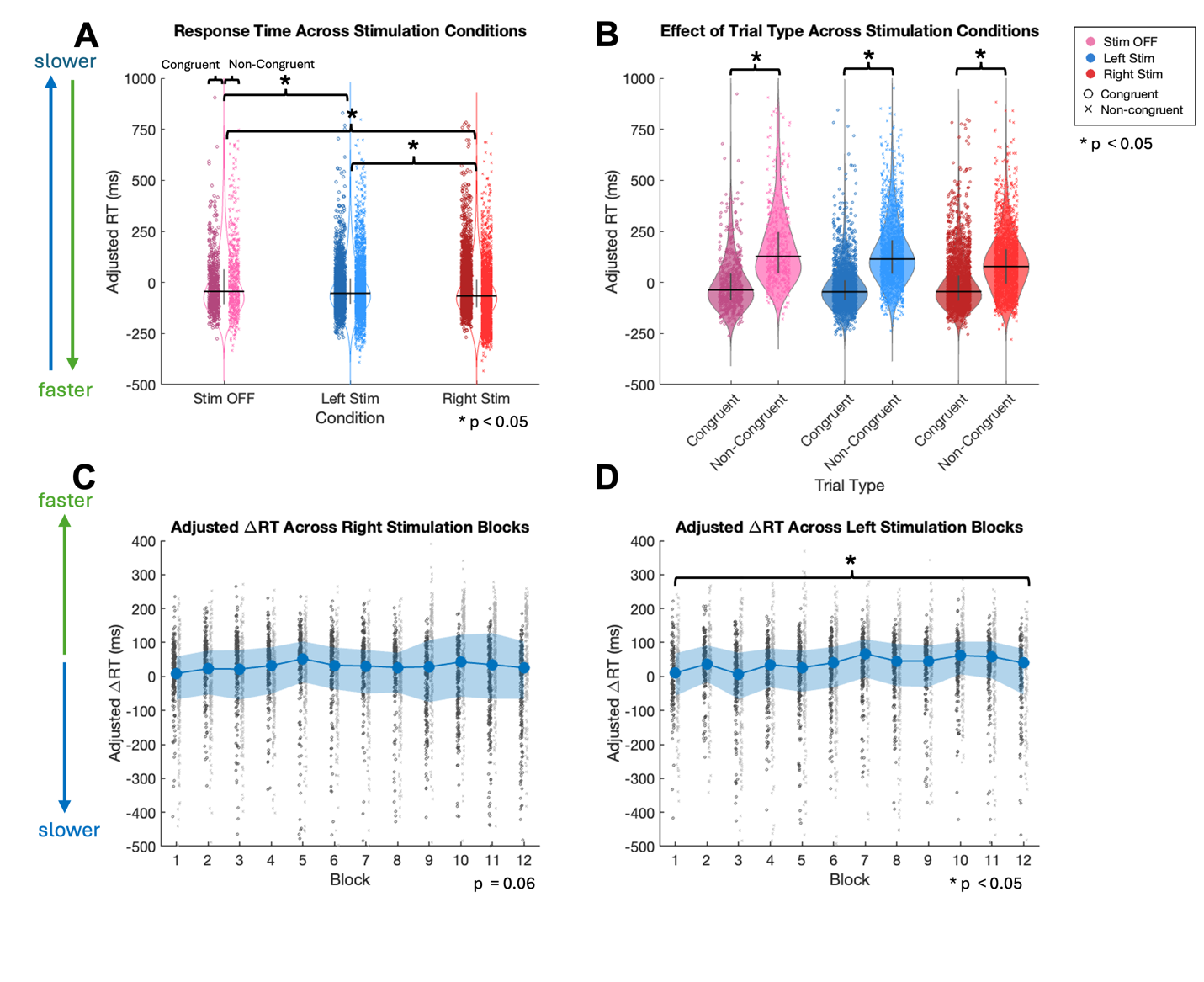

**Supplementary Figure 2.** Complete behavior results with full distribution of data points from the MSIT showing **(A)** effect of stimulation conditions (StimOFF, LeftStim, RightStim) on overall RT, **(B)** effect of trial type (congruent, non-congruent) on overall RT across stimulation conditions, **(C)** effect of right-sided stimulation blocks on ΔRT, and **(D)** effect of left-sided stimulation blocks on ΔRT. Each point corresponds to RT from an individual trial, with dark circles indicating congruent RTs and light crosses indicating non-congruent RTs.

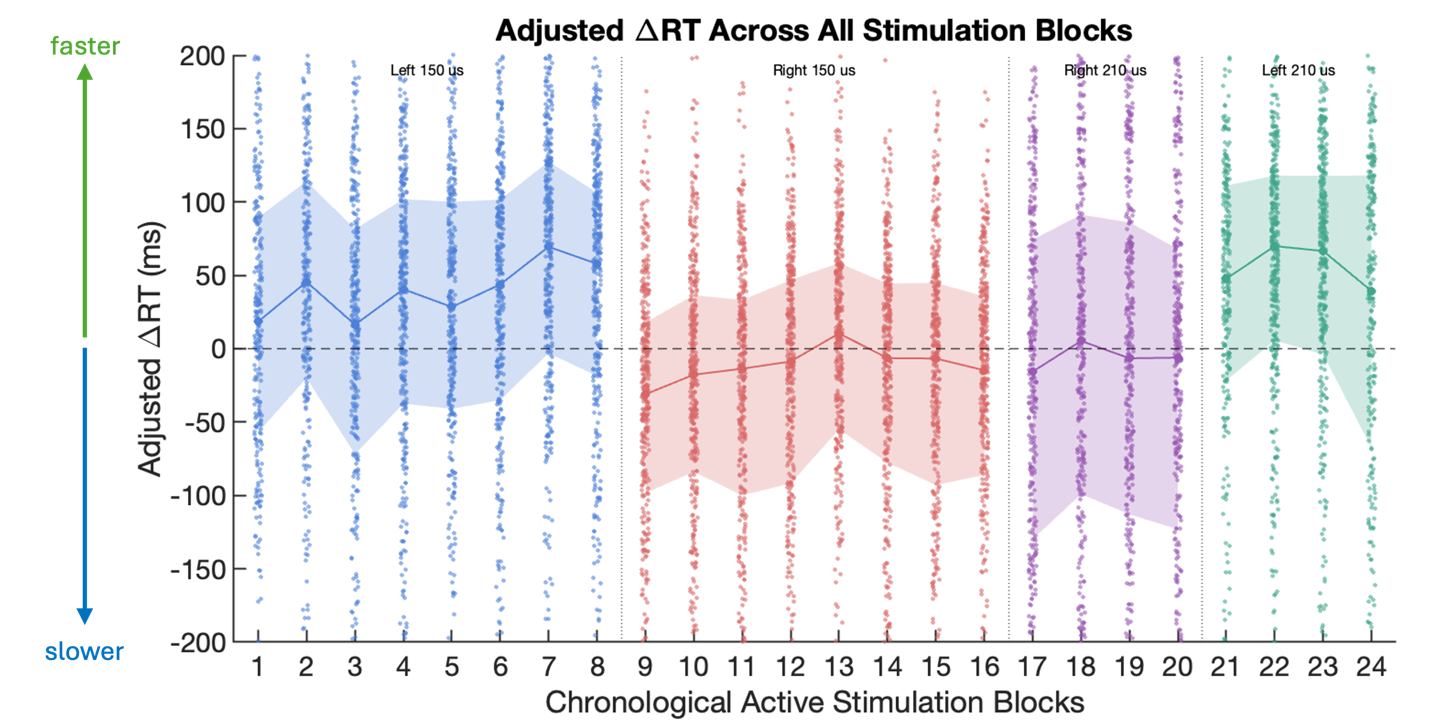

**Supplementary Figure 3.** Adjusted ΔRTs (AdjΔRT) across all 24 active stimulation blocks in chronological order, with median and interquartile range. Each point corresponded to AdjΔRT from an individual trial. A positive value corresponded to a faster AdjΔRT (increased AdjΔRT = faster), and a negative value corresponded to a slower AdjΔRT (decreased AdjΔRT = slower). Blocks 1-8 correspond to left-sided stimulation at 150 µs, blocks 9-16 to right-sided stimulation at 150 µs, blocks 17-20 to right-sided stimulation at 210 µs, and blocks 21-24 to left-sided stimulation at 210 µs.

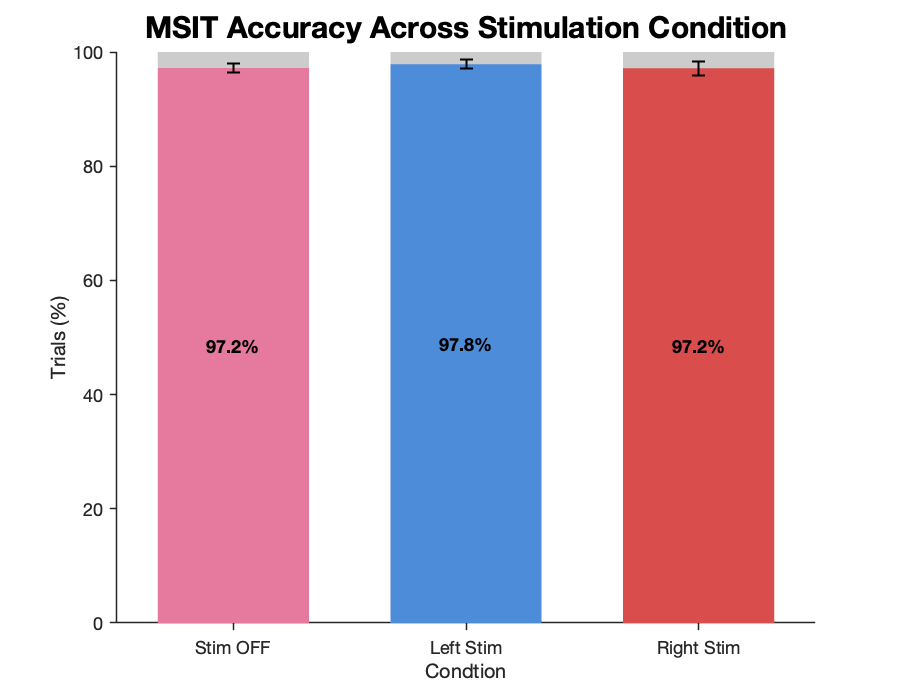

**Supplementary Figure 4.** MSIT accuracy across stimulation conditions. Mean percentage of correct responses during no stimulation (Stim OFF), left-sided stimulation (Left Stim), and right-sided stimulation (Right Stim). Colored bars indicate correct responses, and gray segments indicate incorrect responses. Error bars represent ±1 SEM across participants.

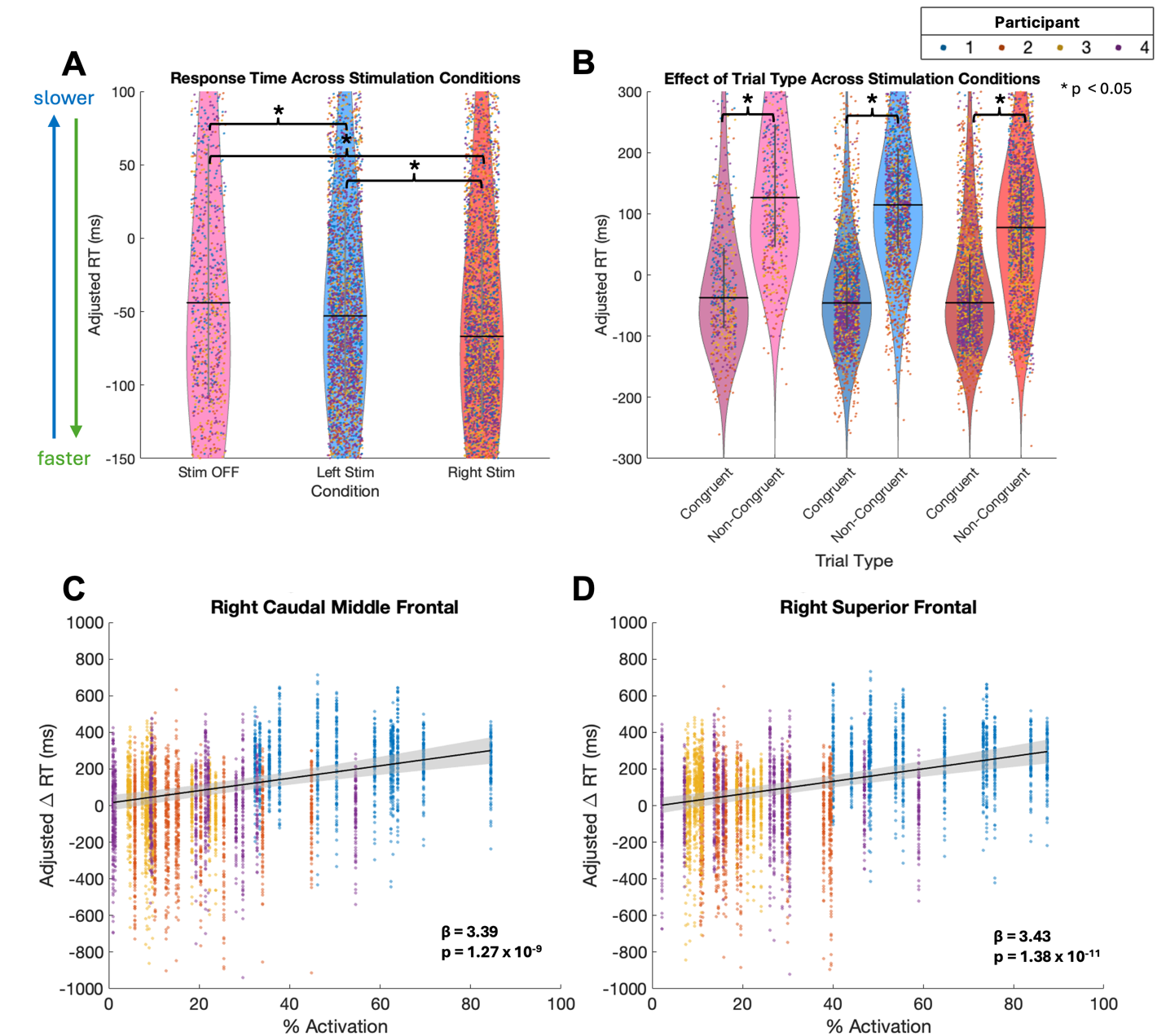

**Supplementary Figure 5. Participant-level MSIT behavioral and pathway activation results. (**A-B) Individual participant adjusted reaction times (AdjRTs) across stimulation conditions (StimOFF, LeftStim, and RightStim) and trial types (congruent and non-congruent). (C-D) Relationships between pathway activation and adjusted change in reaction time (ΔRT) for the right caudal middle frontal (C) and superior frontal (D) pathways. Points are color-coded by participant to visualize individual observations across the four participants; solid lines and shaded regions indicate pooled GLM predictions and 95% confidence intervals, respectively.

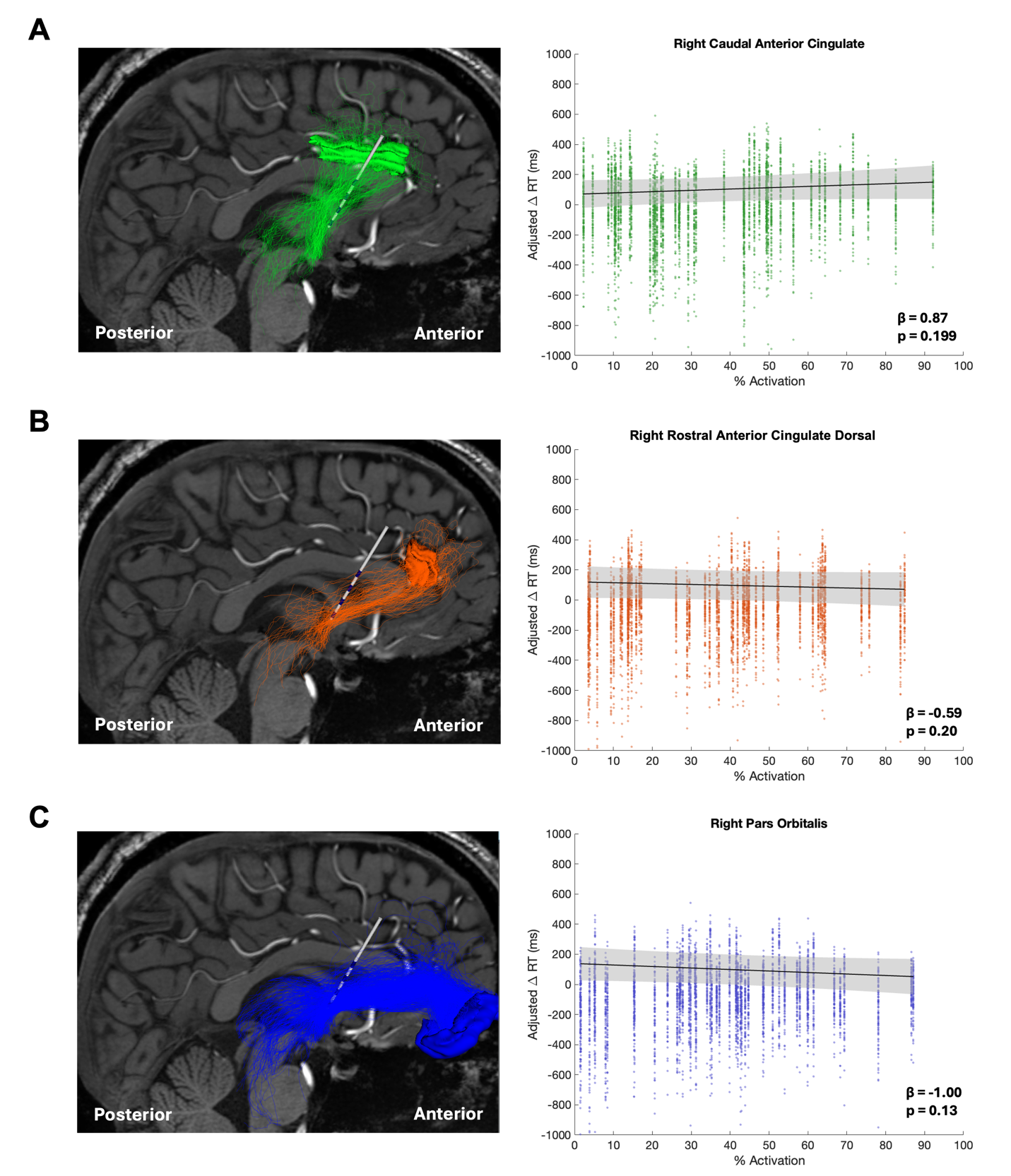

**Supplementary Figure 6.** Pathway activation results for non-significant pathways. Follow-up GLMs assessed the predictive contribution of five pathways identified by the mixed-effects regularized regression and isolated three non-significant pathways. Left subplot displays example pathways in a sagittal orientation from an individual participant, including the implanted DBS lead and its connectivity to its respective PFC region. Right subplot shows the association between pathway-specific percent activation and adjusted ΔRT across all participants with fixed effects prediction from GLM (solid line) and 95% confidence interval (shaded grey). Percent activation of tracts connected to the **(A)** right caudal anterior cingulate gyrus, **(B)** right rostral/dorsal anterior cingulate, and **(C)** right pars orbitalis was neither predictive of a faster nor slower ΔRT. Percent activation values appear discretized due to the use of discrete stimulation amplitudes (2 mA increments), producing step-wise changes in modeled pathway activation.

| **Model** | **Right Pathway AIC** | **Left Pathway AIC** |
| --- | --- | --- |
| ΔRT ~ TrialType + StimConfiguration + (1\|Participant) | 11451.5 | -5317.8 |
| ΔRT ~ TrialType + StimConfiguration + PathwayActivation + (1\|Participant) | 11436 | -5381.3 |
| ΔRT ~ TrialType + PathwayActivation + (1\|Participant) | 11429.7 | -5305.1 |

**Supplementary Table 2.** Summary of mixed-effects models evaluating whether behavioral performance is better explained by stimulation configuration or modeled pathway activation. The best performing model (lowest AIC) was the combination of both in the left hemisphere, and pathway activation alone in the right hemisphere. In neither hemisphere was a model using stimulation configuration superior.

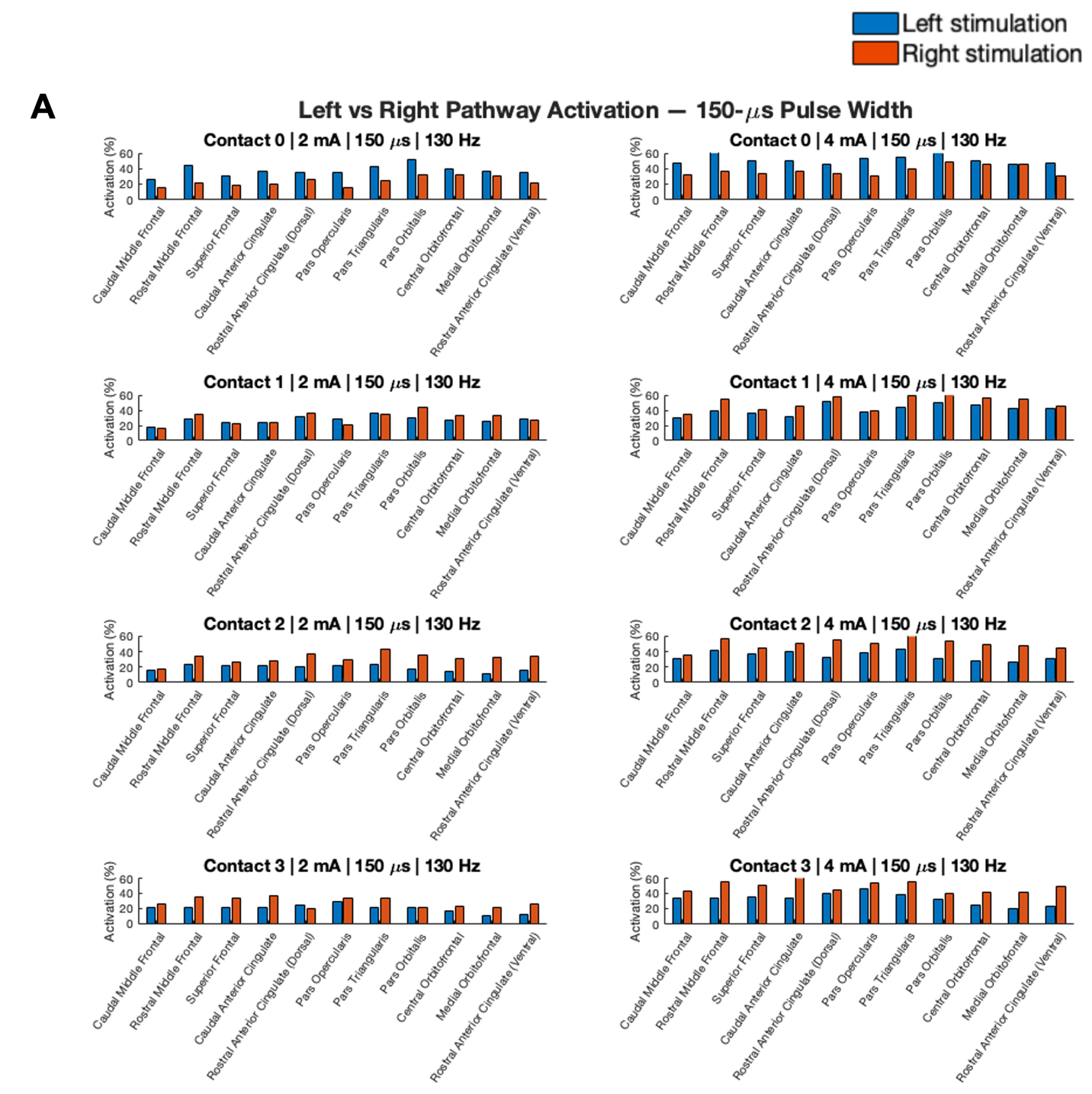

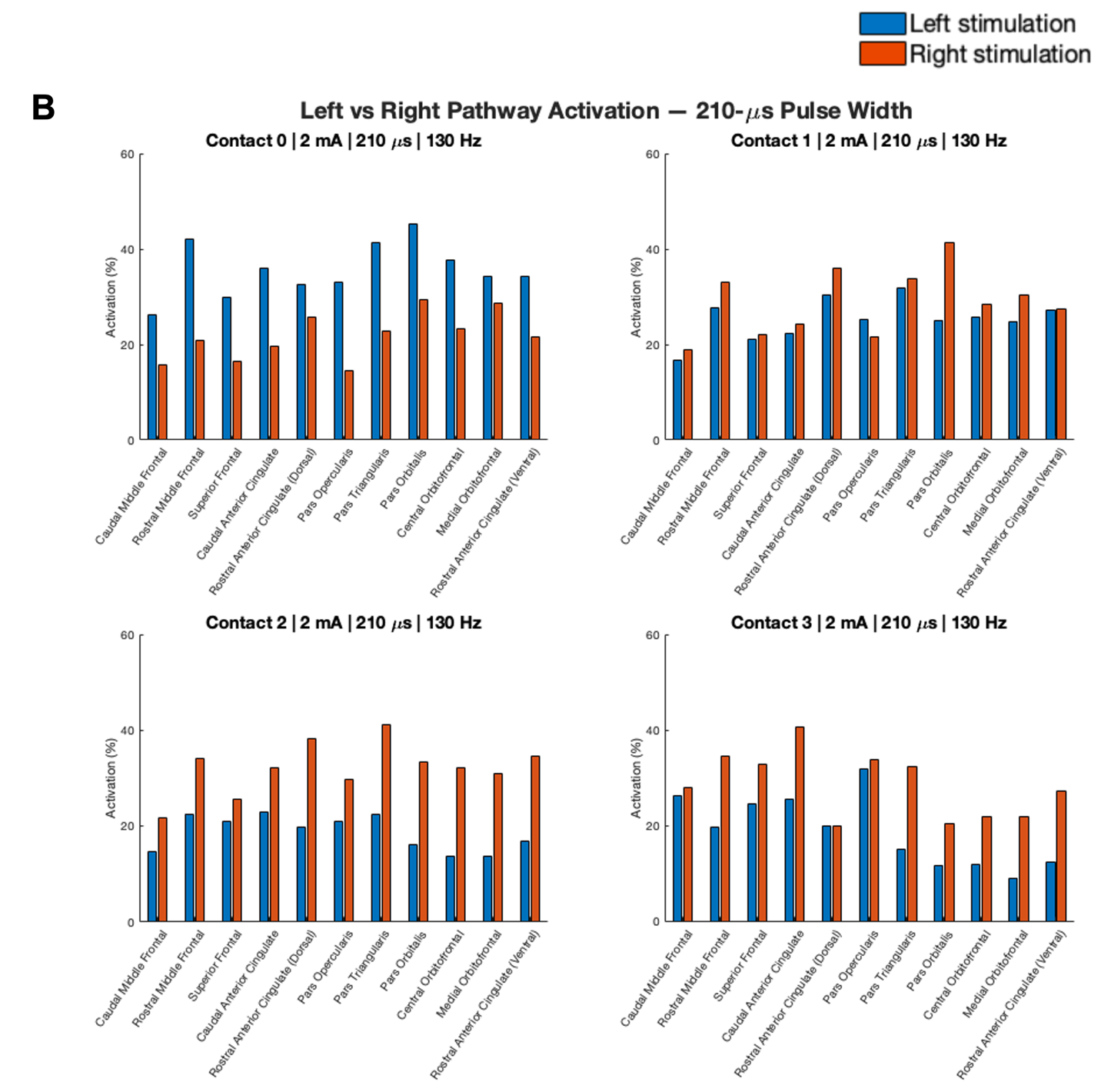

**Supplementary Figure 7.** Hemispheric differences in modeled pathway activation across stimulation configurations. (A) 150 μs and (B) 210 μs pulse width. Bar plots show the mean modeled activation (%) across 11 left-sided ICpre pathways and 11 right-sided ICpre pathways during left-sided (blue) stimulation and right-sided (red) stimulation configurations, respectively. Each subplot represents a stimulation configuration defined by electrode, stimulation amplitude, pulse width, and frequency. Percent activation was averaged across participants within each configuration.
